# HIV-differentiated metabolite N-Acetyl-L-Alanine dysregulates human natural killer cell responses to *Mycobacterium tuberculosis* infection

**DOI:** 10.1101/2023.02.28.530445

**Authors:** Baojun Yang, Tanmoy Mukherjee, Rajesh Radhakrishnan, Padmaja Paidipally, Danish Ansari, Sahana John, Ramakrishna Vankayalapati, Deepak Tripathi, Guohua Yi

## Abstract

**Background:** *Mycobacterium tuberculosis* (*Mtb*) has latently infected over two billion people worldwide (LTBI) and causes 1.8 million deaths each year. Human immunodeficiency virus (HIV) co-infection with Mtb will affect the Mtb progression and increase the risk of developing active tuberculosis by 10-20 times compared to the HIV-LTBI+ patients. It is crucial to understand how HIV can dysregulate immune responses in LTBI+ individuals.

**Methods:** Plasma samples collected from healthy and HIV-infected individuals were investigated by liquid chromatography-mass spectrometry (LC-MS), and the metabolic data were analyzed using an online platform Metabo-Analyst. ELISA, surface and intracellular staining, flow cytometry, quantitative reverse transcription PCR (qRT-PCR) were performed by standard procedure to determine the surface markers, cytokines and other signaling molecule expression. Seahorse extra cellular flux assays were used to measure the mitochondrial oxidative phosphorylation and glycolysis.

**Results:** Six metabolites were significantly less abundant, and two were significantly higher in abundance in HIV+ individuals compared to healthy donors. One of the HIV-upregulated metabolites, N-Acetyl-L-Alanine (ALA), inhibits pro-inflammatory cytokine IFN-□ production by NK cells of LTBI+ individuals. ALA inhibits glycolysis of LTBI+ individuals’ NK cells in response to *Mtb*.

**Conclusions:** Our findings demonstrate that HIV infection enhances plasma ALA levels to inhibit NK cell-mediated immune responses to *Mtb* infection, offering a new understanding of the HIV-*Mtb* interaction and providing the implication of nutrition intervention and therapy for HIV-*Mtb* co-infected patients.

## 1. Introduction

*Mycobacterium tuberculosis* (*Mtb*), the causative pathogen for tuberculosis, is responsible for ∼2 billion of latent infections (LTBI) globally, with 1.8 million deaths each year[1]. Among these *Mtb*-infected individuals, an estimated 13 million are co-infected with human immunodeficiency virus (HIV)[2]. Even though the *Mtb* latent infection is asymptomatic, once co-infected with HIV, the risk of developing active TB in LTBI patients is 10-20 times more compared HIV uninfected LTBI+ individuals [3]. To develop better vaccines and treatment methods, it is important to understand HIV-mediated dysregulation of immune responses in LTBI+ individuals.

Cellular metabolism plays a crucial role in regulating human immune responses to pathogenic infections [4, 5]. The accumulation of specific metabolites from the pathogen-infected cells can function as epigenetic modifiers to the immune cells and alter the epigenetic landscape of some metabolically important enzymes [6], and it leads to changes in immune cell homeostasis and/or functional changes of the affected immune cells [7]. Eventually, it enhances the pathogen’s growth and disease progression. During HIV infection, the metabolism of host cells will be skewed to viral survival and replication[8, 9]. HIV-infected macrophages are known to be metabolically altered with the characteristics of mitochondrial fusion, lipid accumulation, and reduced mitochondrial ATP production[10]. HIV-induced metabolites, such as glucose and some amino acids and their intermediate products, have been reported to significantly impact the function of the immune system. For instance, glucose uptake is essential for the activation of CD4+ T cells and the pro-inflammatory cytokine production in myeloid cells during HIV infection[11–13], mechanistically it regulates CD4+ T cells or myeloid cells via tricarboxylic acid cycle (TCA cycle)[14].

However, little is known about how HIV-driven host metabolite changes can dysregulate the immune system and control *Mtb* immunopathogenesis, which is an important question that needs to be answered in terms of controlling *Mtb* progression in HIV-*Mtb* co-infected patients. In this study, we performed a metabolomics comparison using the plasma from healthy donors and HIV-infected patients (with or without antiretroviral therapy (ART)). We show that one of the metabolites, N-Acetyl-L-Alanine (ALA) is more abundant in HIV+ plasma compared to healthy donor samples. ALA inhibited pro-inflammatory cytokine IFN-γ production by NK cells of LTBI+ individuals. We also found ALA inhibits glycolysis of LTBI+ individuals’ NK cells in response to *Mtb.* Our findings demonstrate that HIV infection enhances plasma ALA levels to inhibit NK cell-mediated immune responses to *Mtb* infection.

## 2. Materials and Methods

### 2.1. Human study sample collection

All healthy (5 samples for each experiment) and HIV-positive plasma samples (8 samples) and LTBI+ blood samples (5-6 samples for each experiment) were collected under the protocols respectively approved by the Institutional Review Boards of Texas Tech University Health Sciences Center at El Paso and the University of University Health Science Center at Tyler. All participants in this study provided written informed consent.

HIV-positive peripheral blood samples were collected into tubes containing sodium heparin and centrifuged at 8,000 × g for 10 min at 4°C for 15 min, and the plasma samples were pipetted out and stored at −80°C until use.

Blood was collected at the Pathology Laboratory of the University of Texas Health Science Center at Tyler, and the PBMCs were isolated using Ficoll-Paque (Fisher Scientific Inc.) density gradient centrifugation as per the manufacturing instructions.

### 2.2 Antibodies and flow cytometry

The antibodies used for this study’s surface and intracellular staining were purchased from Biolegend Inc., CA. These fluorescence-labeled antibodies were used for staining different panels: APC-Cy7-CD3 (clone HIT3a), PE-CD4 (A161A1), PE-Cy7-CD45 (H130), PE-Dazzle 594-CD56 (NCAM) (HCD56), BV605-CD8 (SK1), BV421-FoxP3 (206D), APC-CD25 (M-A251), APC-TNFα (MAb11), BV421-IFN□ (4S.B3), BV510-KLRG1 (2F1/KLRG1), APC-CD27 (M-T271), FITC-CD4 (SK3), BV711-CD25 (BC96), PerCP-Cy5.5-PD-1 (EH12), PE-Cy7-CD8 (SK1), BV421-CCR7 (G043H7), BV605-CD56 (HCD56), PE-CD62L (DREG-56), PE-Cy5-CD19 (HIB19), BV711-CD16 (3G8), FITC-CD14 (HCD14), BV421-CD14 (HCD14), BV605-CD11c (3.9), BV605-CD11b (ICRF44), PB-CD45 (2D1), and APC-CD40L (24-31). The isotype antibodies used for this study are (same clones were chosen as the above fluorescence-labeled antibodies): APC mouse IgG1, BV421 mouse IgG1, PerCP/Cy5.5 mouse IgG1, BV605 mouse IgG1, PE/Dazzle 594 mouse IgG1, BV711 mouse IgG1, PE Rat IgG2b, BV510 mouse IgG2a, APC mouse IgG1, APC-Cy7 mouse IgG1, PE-Cy7 mouse IgG1, FITC mouse IgG1, and PB mouse IgG1. For the surface staining, the cells were stained by different panels of fluorescence-labeled antibodies for 30 min on ice, and the stained cells were then washed in FACS buffer (2% fetal calf serum (FCS) in PBS), and resuspended in 500 µl FACS buffer. For intracellular staining, the surface-stained cells were fixed for 30 min at room temperature and intracellularly stained with BV421-IFN □ (4S.B3) in 1x permeabilization buffer for 20 min in room temperature using Intracellular Fixation & Permeabilization Buffer Set (eBioscience™; 88-8824-00). The stained cells were collected by Attune NXT (Thermo Fisher Scientific), and the data were analyzed with FlowJo (Tree Star, Ashland, OR, USA). Dead cells were removed by both forward and side scatter gating.

### 2.3. PBMC treatments with various metabolites

Two million of PBMCs each well in 12-well plate were stimulated with 10 µg/ml □-irradiated *Mtb* (□ *Mtb*) and immediately followed by treatments with different concentrations of various metabolites. The unstimulated cells and stimulated but untreated cells served as negative controls. After 72 h, the cells were collected for surface and intracellular staining, and the supernatants were collected for ELISA to test the cytokine expressions.

### 2.4. Quantitative reverse transcription PCR (qRT-PCR)

*The* □*Mtb*-stimulated, ALA-treated and untreated PBMCs were collected, and the RNA was extracted by using TRIzol reagent (Invitrogen) as recommended by the manufacturer.

The mRNA transcription levels of the NK cell signaling molecules and the death pathways’ molecules were measured by qRT-PCR using βActin as an internal control with specific primer sets (Integrated DNA Technologies) (see Table S1 and S2 in the supplemental material).

### 2.5. Extracellular flux measurement

PBMC from LTBI+ healthy donors were plated in 12 well plates at a concentration of ∼5×10^6^ cells/ well. Cells were treated with either N-acetyl-L-alanine or □*Mtb* or both, along with no treatment for up to 48 hours. After 48 hours, NK cells were isolated from respective wells using NK cell isolation kit (Miltenyi biotec; Cat: 130-092-657) following standard protocol and plated at a concentration of 2×10^5^ cells per well of a seahorse XFe96 assay plate in seahorse XF DMEM media (Agilent; 103575-100) supplemented with 1mM pyruvate, 2mM glutamine and 10mM of glucose. OXPHOS measurements were performed in a Seahorse Xfe96 Analyzer, using mito-stress test kit (Agilent; 103015-100). Measurement of OCR (oxygen consumption rate) was done after the subsequent addition of 1.5 μM oligomycin, 1μM FCCP (Carbonyl cyanide-4 trifluoromethoxy phenylhydrazone) and 0.5 μM Rotenone/Antimycin A(Rot/AA). Basal respiration is measured as OCR after subtracting non mitochondrial respiration obtained after adding Rot/AA, spare respiratory capacity is measured as the highest respiration obtained compared to basal after adding FCCP, ATP coupled respiration is amount of OCR affected by the addition of Oligomycin. Glycolytic parameters were measured using the glycolysis stress test kit (Agilent; 103020-100), measurement of ECAR (extracellular acidification rate) was done after the sequential addition of 1mM Glucose, 1.5 μ M Oligomycin, 5mM 2-deoxyglucose(2-DG). Basal glycolysis was measured as the resting ECAR value after the addition of glucose, glycolytic capacity is defined as the maximum ECAR after the addition of oligomycin, and glycolytic reserve is the difference between basal and maximal glycolytic capacity. Wave Desktop 2.6 software (Agilent) was used for the data analysis.

### 2.6. Metabolomics

Plasma samples were collected from healthy and HIV-infected individuals at Texas Tech University Health Science Center El Paso. Plasma samples were analyzed at the metabolomics core facility at the Children’s Medical Center Research Institute at UT Southwestern (Dallas, TX) using liquid chromatography-mass spectrometry (LC-MS). A triple-quadrupole mass spectrometer was used in MRM mode for the analysis, with two different dilutions for the samples, including four different retention times and three quality control samples. Further annotation of peaks was done using a proprietary database. The data matrix was statistically arranged using Metabo-Analyst (https://metaboanalyst.ca), an online platform for reading metabolomics data using default parameters.

### 2.7. ELISA and LDH assay

All ELISA kits were purchased from ThermoFisher Scientific Inc, CA. The supernatants were collected from the PBMC culture after 72 h treatments. The ELISA procedures to detect the IFN-□, TNF-α, IL-1β, IL-4, IL-13 and IL-17A were performed according to the manufacturer’s protocols. A colorimetric CyQUANT lactate dehydrogenase (LDH) assay (ThermoFisher Scientific Inc.) was performed to determine the LDH activity in culture supernatants of PBMCs.

### 2.8. Statistic analysis

Each treatment was triplicated (qRT-PCR) or duplicated (all other experiments), and the experiments were repeated at least once to ensure reproducibility. Power analysis was done to determine the sample size to ensure biological significance. The data were analyzed by GraphPad Prism software. Paired student T-test was used to analyze the difference between the treated and untreated samples from the same donor. Statistical significance was defined as *P≤0.05, **P≤0.01, and ***P≤0.001.

## 3. Results

### 3.1. Metabolic profiles of HIV-positive patient plasma

To characterize the HIV patient-specific metabolic landscape, we performed liquid chromatography-mass spectrometry (LC-MS) based metabolic profiling of plasma samples from HIV-positive patients (both treatment-naive and ART-treated) and healthy donors. We performed supervised partial least squares discriminant analysis (PLS-DA) of metabolome profiles and plotted the two principal components explaining the highest magnitude in Figure 1a. We observed that the plasma metabolome landscape of HIV patients (Both treatment naïve and ART-treated) is distinct from healthy control, while profiles of treatment naïve and ART-treated patients were similar (Figure 1a). To identify metabolites that are altered in HIV infection, we performed differential enrichment analysis and identified 60 metabolites that showed altered abundance in patients compared to healthy donors at a false discovery rate (FDR) <0.05. We performed hierarchical clustering on differentially enriched metabolites (Figure 1b). As expected, healthy donors and HIV patients formed distinct clusters which is consistent with the PLS-DA results (Fig 1b).

**Figure 1.**
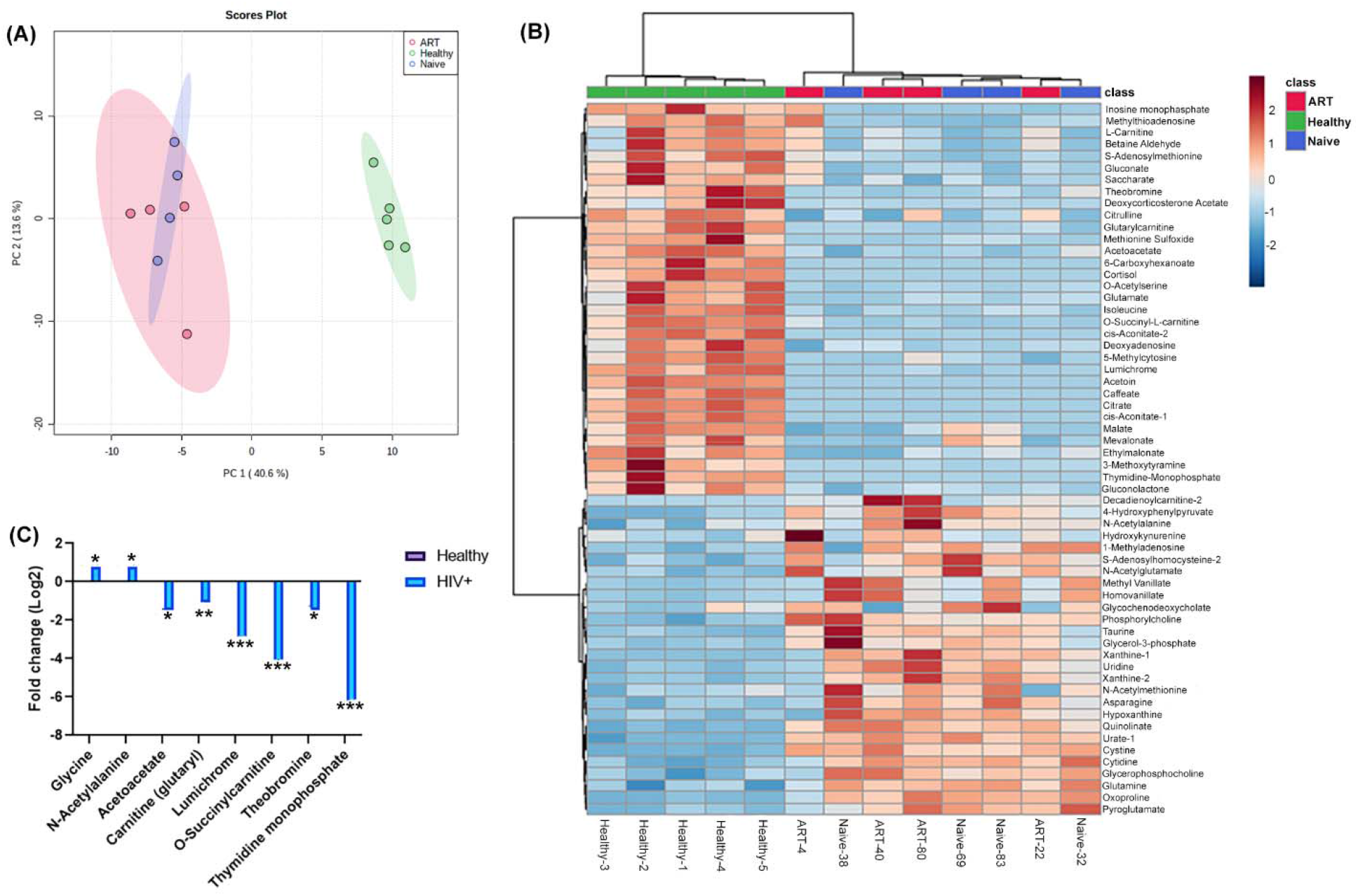
Plasma metabolic profile of HIV+ patients (treatment naïve or ART-treated). **(A)** Scatter plot showing partial least squares discriminant analysis (PLS-DA) of plasma metabolomic profiles. The two principal components explaining highest variance were plotted on X and Y axis. **(B)** Heatmap shows the top 60 differentially abundant metabolites identified in a comparison between healthy donors and HIV patients at 5% FDR. **(C)** Fold change in abundance of 6 selected metabolites computed from metabolome profiles in a comparison between HIV patients and healthy donors. The bars represent log2 fold change in HIV patients compared to healthy donors. The asterisks *, **, and *** denote FDR<0.05, FDR<0.01, and FDR<0.001, respectively.

However, treatment naïve and ART-treated patients formed a single cluster underscoring similarities in the metabolome profiles of both treatment groups. The most significant metabolites altered between healthy donors and patients were selected using the following criteria: 1) the FDR value ranks beyond the first 60; 2) the VIP score is >1 by PLS-DA analysis; and 3) The fold change is >1.5 (HIV/healthy donors for upregulated metabolite, and healthy/HIV+ donors for down-regulated metabolites). Finally, we found that there were two metabolites, N-Acetyl-L-alanine (ALA) and glycine, were upregulated in HIV-positive plasma samples, and six metabolites were downregulated, they are acetoacetate, glutarylcarnitine, lumichrome, O-Succinylcarnitine, theodromine, and thymidine monophosphate, respectively (Figure 1c).

### 3.2. ALA inhibits IFN-□ and TNF-α secretion by -irradiated Mtb (Mtb) stimulated

Pro-inflammatory cytokines IFN-□, TNF-α, IL-17A, and IL-1β are known to play important roles in controlling *Mtb* infection[15] and anti-inflammatory cytokine IL-10 play an important role in the reactivation of TB [16], We asked whether the above identified metabolites have any effect on these cytokines production by PBMC obtained from LTBI+ donors. As shown in Fig. S1, we performed LDH assay to select the optimal concentrations of the metabolites for *in vitro* experiments. We cultured PBMC from healthy LTBI+ donors with or without □*Mtb,* as mentioned in the methods section. Some of the □*Mtb* cultured PBMC were cultured with the metabolites. After 72 h, culture supernatants were collected, and cytokine levels were determined by ELISA. ALA (3 μM concentration) significantly inhibited □*Mtb* stimulated IFN-□, TNF-α and IL-17 production by PBMCs of LTBI+ individuals (Fig. 2a, b and c). In contrast, other metabolites had no effects on IFN-□, TNF-α and IL-17 production of LTBI+ PBMCs (Fig. 2a, b and c). None of the metabolites have any effect on IL-1β, IL-10 and IL-13 production(Fig. 2d, e and f).

**Figure 2.**
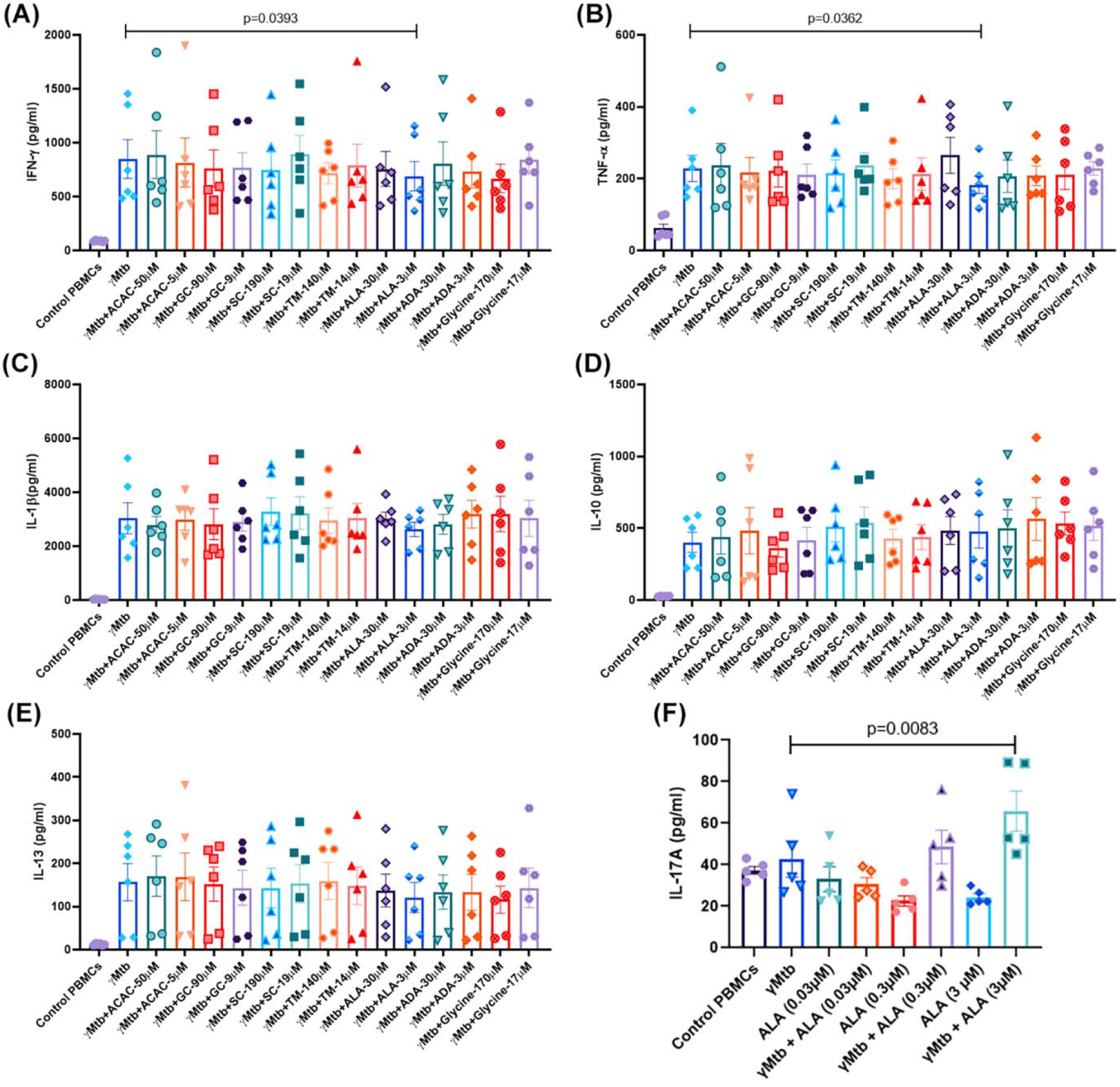
Effect of metabolites on cytokine production by PBMC of LTBI+ donors. PBMC of healthy LTBI+ donors were cultured with or without γ*Mtb* (10 μg/mL) and in g/mL) and in the presence or absence of different concentrations of ALA and other metabolites as mentioned in the methods section. After 72 hours, culture supernatants were collected, and cytokine levels were measured by ELISA. **(A)** IFN- □,, **(B)** TNF-α, **(C)** IL-1β, **(D)** IL-10, **(E)** IL-13, and **(F)** IL-17A. In (A) to (E), six donors were collected, while in (F), five donors were collected. Paired T-tests were used to compare the differences between untreated and treated PBMCs from the same samples. The mean values and SDs are shown, and the significant P values are shown (p<0.05).

### 3.3. ALA inhibits IFN-□ secretion of NK cells

We determined the effects of ALA on the expansion of various immune cell populations in the above-cultured cells. We found that ALA triggered the expansion of the classic monocytes (CD14+CD16-) population, while it did not affect the expansion of other immune cells and their subpopulations (Fig. S2). To determine the cellular source for the IFN-□ and TNF-α, various cell populations in the above-cultured cells were sorted (cell purity is shown in Fig. S3), and a quantitative RT-PCR (qRT-PCR) was performed to determine the IFN-□ and TNF-α transcription levels. ALA significantly inhibited IFN-□ gene expression by NK cells and CD8+ T cells (Fig. 3a). In contrast, TNF-α gene expression by the above immune cell populations was not affected by ALA (Fig. 3b). We further confirmed the above findings at the protein level by performing intracellular staining on the above immune cell populations and found that ALA inhibits IFN-□ production by NK cells in response to □*Mtb* (Fig. 4c and d). In contrast, ALA had no effect on IFN-□ production by CD8+ and CD4+ cells (Fig. 4a, b, and d).

**Figure 3.**
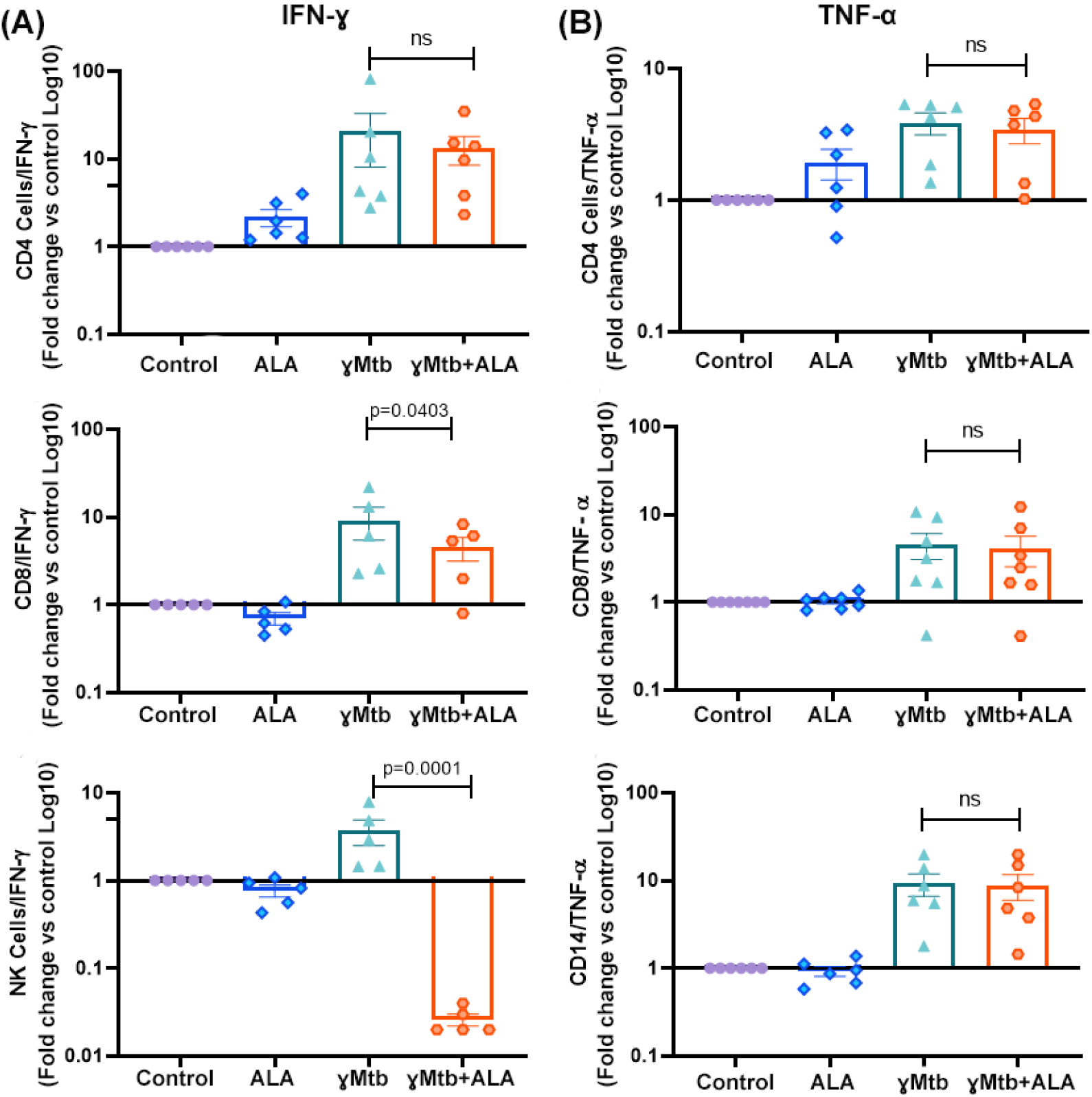
IFN-□ and TNF-α gene expression profile of various immune cells in □*Mtb* cultured PBMC of LTBI+ donors. PBMCs of healthy LTBI+ donors were cultured with or without γ*Mtb* (10 μg/mL) and in the presence or absence of ALA as mentioned in the methods section. After 48 hours, various immune cells were isolated by flow sorting, RNA was collected, and real-time PCR analysis was performed to determine IFN-□ and TNF-α gene expression. **(A)** IFN- L transcription level in different cell types. (**B)** TNF-α transcription level in different cell types. PBMCs from five LTBI+ donors were used for the study. Paired T-tests were used to compare the differences between untreated and treated PBMCs from the same samples. The significant P values are shown (p<0.05).

**Figure 4.**
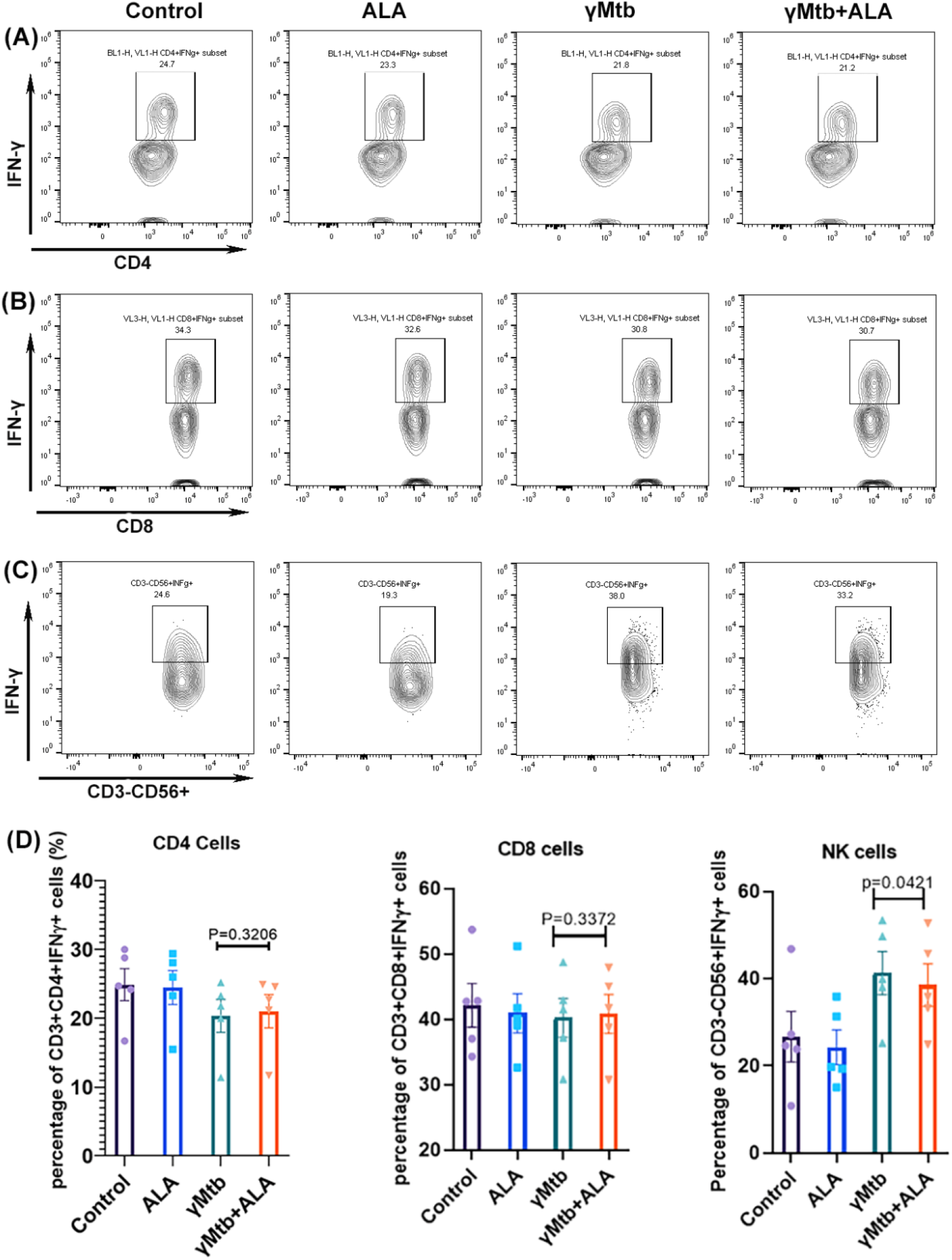
IFN-□ levels of various immune cells in □ *Mtb* cultured PBMC of LTBI+ donors. PBMCs of healthy LTBI+ donors were cultured with or without γ*Mtb* (10 μg/mL) and in the presence or absence of ALA as mentioned in methods section. After 48 hours, intracellular staining was performed to determine IFN-□ levels of various immune cell populations. **(A, B, and C)** IFN- L staining of a representative donor PBMCs, the top, middle and bottom panels represent CD4+ T cells (A), CD8+ T cells (B), and NK cells (C), respectively. The percentages of IFN- L positive cells are shown. **(D)** Collective summary of IFN- positive CD4+, CD8+, and CD56+ cells of five donors. L Paired T-tests were used to compare the differences between untreated and treated PBMCs from the same samples. The significant P values are shown in the figure.

### 3.4. ALA inhibits nuclear factor kappa B (NF-□B), activator protein-1 (AP1), and antimicrobial peptides expression by □Mtb cultured NK cells

We cultured PBMC from healthy LTBI+ donors with or without □*Mtb* stimulation as mentioned in the methods section. Some of the □*Mtb* cultured PBMCs were cultured with 3 μM ALA. After 48 h, NK cells were sorted, and qPCR was performed to determine the expression of 22 transcription factors and signaling molecules. Among these ALA significantly inhibited NF-□B, AP1 and antimicrobial peptides GZMA and GZMB gene expression by □*Mtb* cultured NK cells. In contrast, SATA4 expression by □*Mtb* cultured NK cells was significantly upregulated by ALA.

### 3.5. ALA did not alter the cell death molecules in □Mtb cultured NK cells

In the above-cultured NK cells, we also determined the expression of 11 key genes involved in various death pathways (i.e., autophagy, apoptosis, pyroptosis, necroptosis, and ferroptosis). We found that ALA did not affect the cell death pathways tested in this study, while Atg3 expression (autophagy-related gene) alone was significantly upregulated when compared to non-treated and γ*Mtb-*stimulated cells (p=0.0023) (Fig. 6).

**Figure 5.**
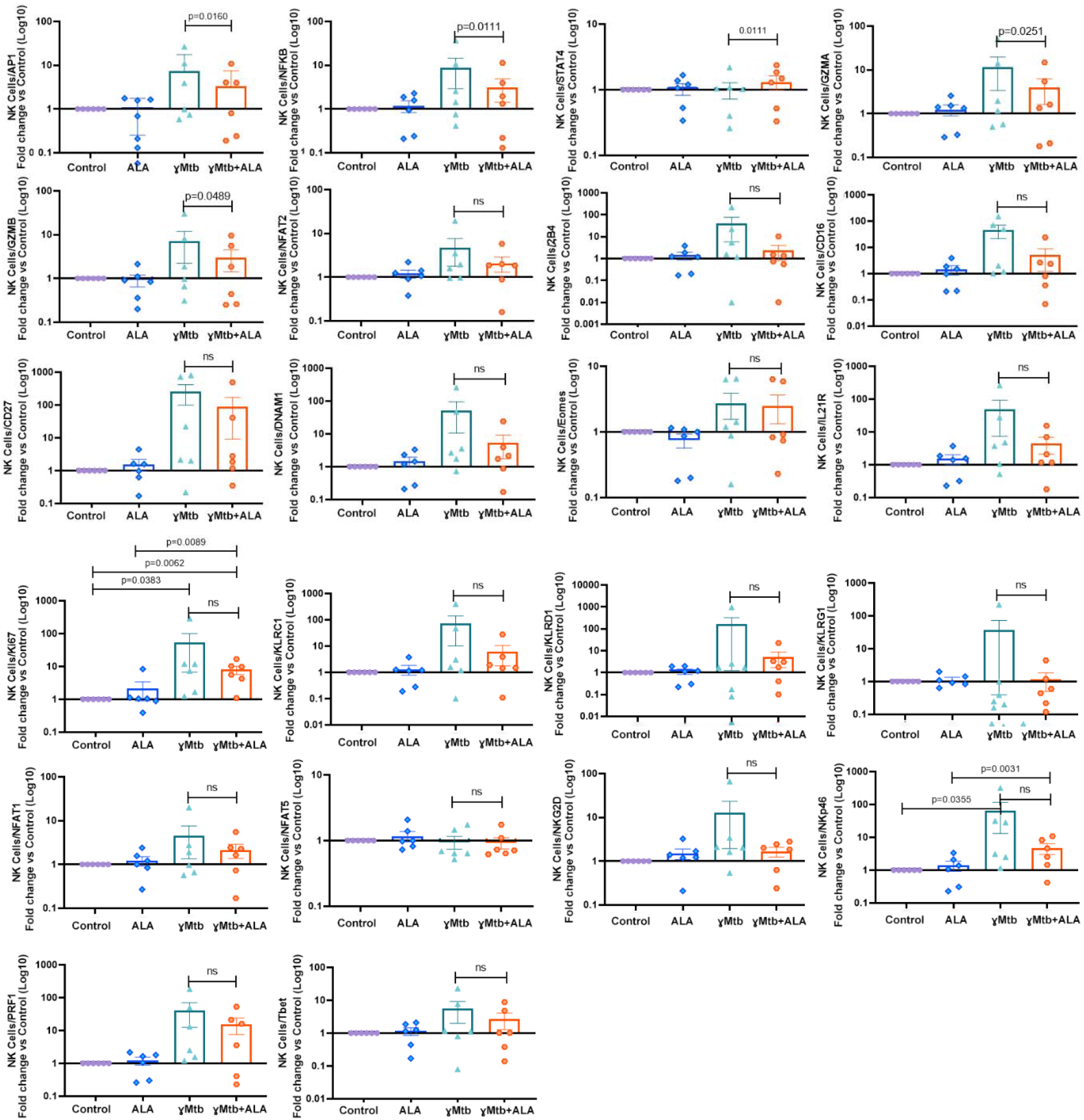
ALA inhibits NF- B, AP1, GZMA, and GZMB gene expression by γ*Mtb* cultured NK cells. PBMCs of healthy LTBI+ donors were cultured with or without γ g/mL) and in the presence or absence of ALA as mentioned in the methods section. After 48 hours, NK cells were isolated by flow sorting, RNA was collected, and real-time PCR analysis was performed to determine various signaling molecules and transcription factors. PBMCs from six LTBI+ donors were used for this experiment. Paired T-tests were used to compare the differences between untreated and treated PBMCs from the same samples. The significant P values are shown (p<0.05). ns: not significant, p>0.05.

**Figure 6.**
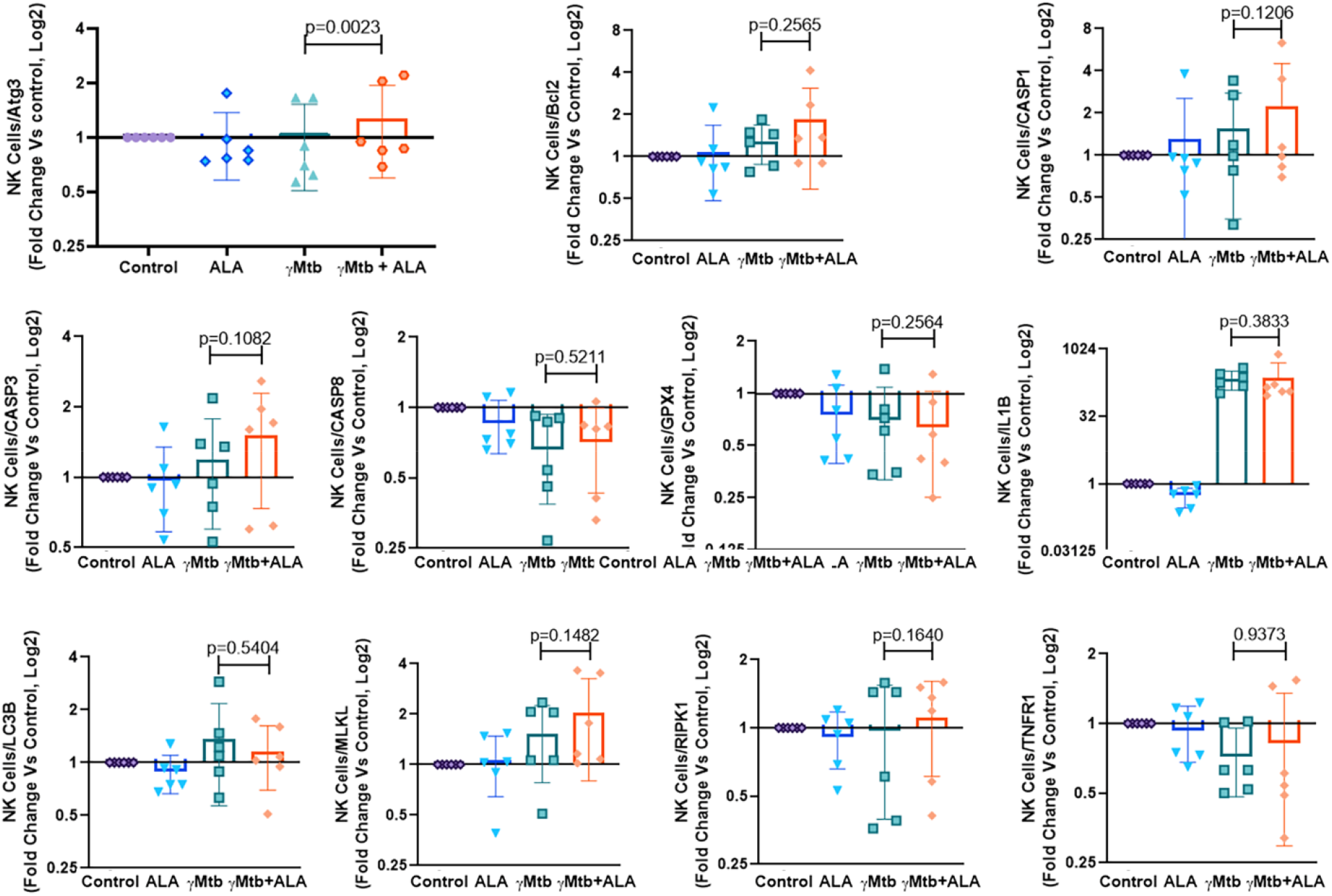
ALA inhibits antimicrobial peptide expression and enhances Atg3 expression by γ*Mtb* cultured NK cells. PBMCs of healthy LTBI+ donors were cultured γ g/mL) and in the presence or absence of ALA as mentioned in μ the methods section. After 48 hours, NK cells were isolated by flow sorting, RNA was collected, and real-time PCR analysis was performed to determine various death pathway gene expressions. PBMCs from six LTBI+ donors were used for this experiment. Paired T-tests were used to compare the differences between untreated and treated PBMCs from the same samples. The P values are shown in the figure.

### 3.6. ALA restricts the bioenergetic machinery in NK cells

Metabolic switch to a glycolytic/energetic phenotype supports diverse NK-cell functions [17, 18]. We determined whether ALA treatment affects the metabolic state of γ*Mtb*-cultured NK cells. We performed a metabolic flux assay (as mentioned in the Methods section) to detect changes in the mitochondrial oxygen consumption rate (OCR) and rate of extracellular acidification (ECAR) as measures of oxidative phosphorylation and glycolysis, respectively.

Freshly isolated PBMCs from LTBI donors (n = 3) were cultured in the presence of γMtb. Some of the γMtb cultured PBMCs were also supplemented with the ALA (3 μM). After 48h, NK cells were isolated from the cultured PBMCs and metabolic flux assay was performed using a seahorse analyzer as mentioned in the methods section. As shown in Fig. 7a and b, various parameters such as basal respiration, ATP production, and spare respiratory capacity of oxidative phosphorylation were significantly reduced in the NK cells from ALA-alone treated PBMCs and γMtb alone cultured PBMCs than control PBMCs. However, we observed a significant marginal reduction in the basal ATP production rate in the NK cells from the PBMCs cultured with γMtb and ALA together than γMtb alone. Surprisingly, we saw pronounced changes in glycolytic parameters as well in the NK cells from ALA alone treated PBMCs and γMtb alone cultured PBMCs compared to control PBMCs (Fig. 7c and d). Interestingly, we found ALA treatment further significantly reduced basal glycolysis, glycolytic capacity, and glycolytic reserve in NK cells from □Mtb cultured PBMCs than γMtb alone cultured PBMCs (Fig.7b). Herein, we observed that ALA treatment significantly suppressed OXPHOS and glycolysis in NK cells, which suggest a higher level of ALA in HIV patients could induce quiescent phenotype in NK cells.

**Figure 7.**
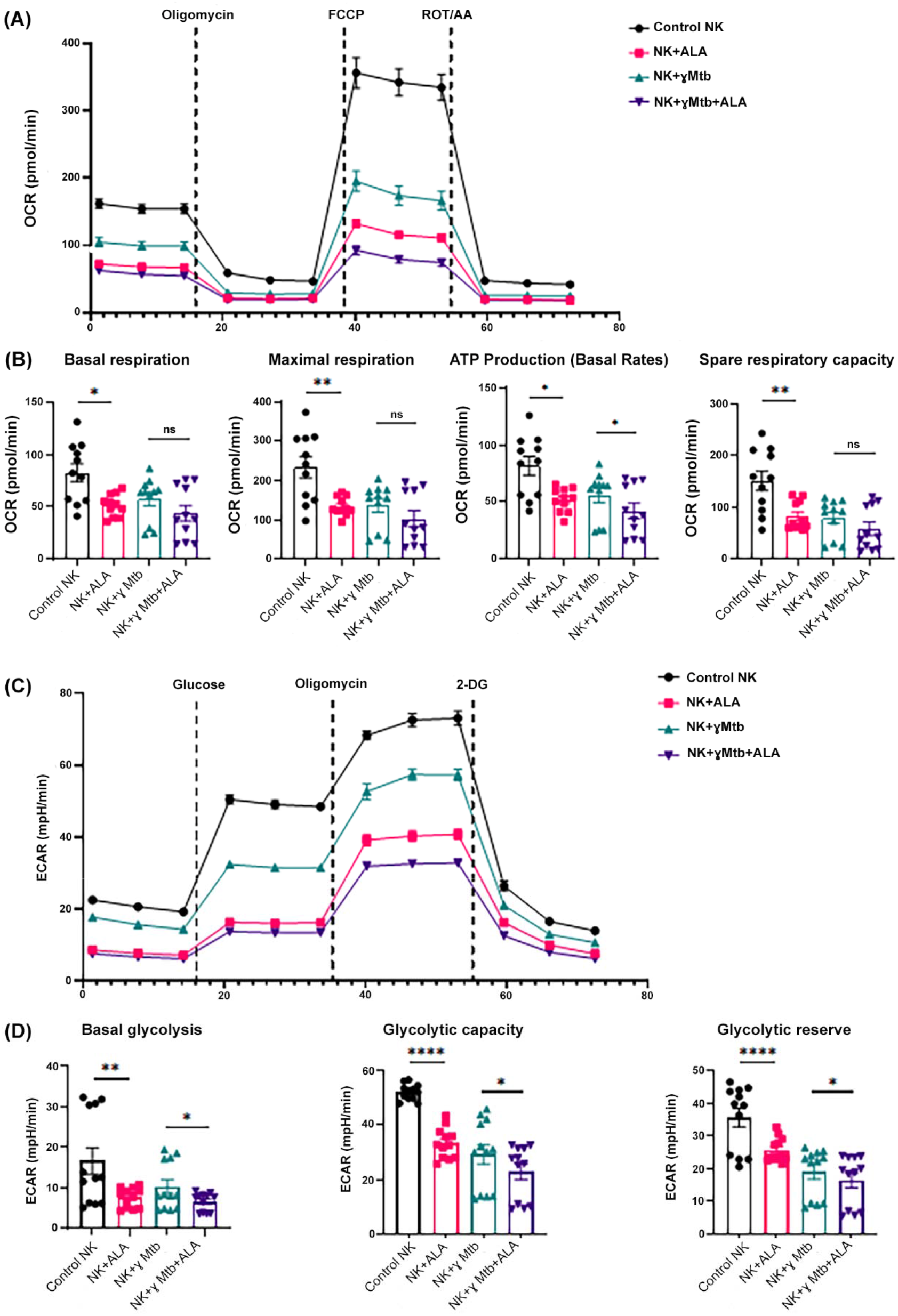
ALA treatment switches NK cells to an energetically quiescent state. PBMCs of healthy LTBI+ donors were cultured with or without γ*Mtb* (10 μg/mL) and in the presence or absence of ALA (3 µM), as mentioned in the methods section. After 48 hours, NK cells were isolated by magnetic cell sorting and subjected to extracellular flux analysis using an Agilent Seahorse XFe96 analyzer. NK cell glycolysis was measured with the sequential addition of Glucose, Oligomycin, and 2-DG. Similarly, OXPHOS parameters were measured in isolated NK cells after the addition of Oligomycin, FCCP and Rotenone/antimycin. **(A and B)** mitochondrial OCR and **(C and D)** ECAR were measured. (A) and (C) respectively show mitochondrial OCR (A) and (C) ECAR in real time as kinetic graphs. (B) shows the collective OXPHOS parameters of Basal respiration, Maximal respiration, ATP production, and Spare respiratory capacity as bar graphs. The P values were derived using an unpaired 2-tailed independent t-test. The mean values and SEMs are shown. (D) Bar graphs show the collective glycolytic parameters such as Basal glycolysis, glycolytic capacity, and glycolytic reserve. In B and D, for all panels, the data is presented as Mean ± SEM (n=12; 4 statistical replicates from 3 individual donors), each parameter between treatments was compared using independent Student’s t-test: *P < 0.05, **P < 0.01, ****P < 0.0001.

## 4. Discussion

Immunometabolism plays a central role in host-*Mtb* interactions and controls the infection outcomes [19–21]. It is not known whether metabolic changes during HIV infection alter immune responses to *Mtb* infection. In the current study, we found HIV infection alters the plasma metabolic profile. Among various elevated metabolites, ALA significantly inhibited NK cell-mediated immune responses to *Mtb* infection. We also found ALA inhibits the expression of two transcription factors NF-□ GZMB, which are important in the activation and antimicrobial activity of NK cells.

Activated NK cells produce IFN-□, which activates macrophages to kill intracellular organisms [22–24]. It has been demonstrated that human NK cells have the potential to contribute to both innate and adaptive immune responses to *Mtb* [25–30]. NK cells can lyse *Mtb*-infected monocytes and alveolar macrophages through the NKp46 receptor and NKG2D [26, 29], and NK cells contribute to the capacity of CD8+ T-cells to produce IFN-□ and to lyse *Mtb*-infected monocytes [27]. During chronic HIV infection, an abnormal, dysfunctional CD56neg NK cell subset expands, and potentially protective NK cell responses are depressed [31–33]. However, limited information is available on the NK cell response to pathogens, including *Mtb*, especially in HIV+LTBI+ individuals.

HIV infection induces significant immunometabolic changes in the host [34–36]. NK cell cytotoxicity and cytokine production depend on their metabolism [37], and altered metabolism is linked to NK cell dysfunction [38]. No information is available about the metabolic requirements of NK cells during *Mtb* and/or HIV infection. Metabolomics provides a versatile tool to study the host immune responses to pathogen infections because metabolism offers a source of energy required for immune cell function. During HIV and *Mtb* infection/co-infection, immune cell activation and inflammation have also been correlated to metabolic changes in the immune cells [39–42], However, the mechanisms how specific metabolites affect the immune responses to HIV/*Mtb* infections remain elusive. Our study shows that HIV-induced metabolite ALA can inhibit IFN-□ production and antimicrobial peptide expression by NK cells and inhibits NK cell glycolysis in response to *Mtb*. As glycolysis is essential to maintain cell viability and inflammation activity [43, 44], the depressed glycolysis, therefore, caused NK cell autophagy and the reduction of IFN- production, as demonstrated in this study. IFN-□ is one of the major cytokines that can limit *Mtb* growth, thus, increased ALA levels during HIV infection can enhance *Mtb* growth and disease progression. It is worth noting that we identified the commonly important metabolites between the treatment naïve and ART-treated samples when performing the metabolomics analysis. This strategy may help discover the potential nutrition intervention/therapy that is suitable for both ART-treated and non-treated HIV-*Mtb* co-infected patients.

Amino acids participate in energy production during cell metabolism, and some amino acids are involved in oxidative stress and redox signaling during HIV and *Mtb* infection[39, 40]. These physiological activities are tightly coupled with immune activation, as evidenced by the study of alanine is essential for CD4+ T cell activation[45]. ALA is the derivative of L-alanine. It can be produced by direct synthesis of N-acetyltransferases or via proteolytic degradation of N-acetylated proteins by hydrolases, such as Aminoacylase I[46]. In our study, the ALA upregulation suggests a decrease in the non-acetylated L-alanine. Consequently, the redox-sensitive transcription factors such as NF-□ b and AP-1 will be downregulated via the redox signaling pathway, and this is also the case in the LTBI NK cells (Fig. 5). In another scenario, ALA may be able to compete with non-acetylated L-alanine to bind to the same nutrition receptor, leading to the decrease of the L-alanine uptake and other similar consequences.

NK cells have been reported to use glycolysis and oxidative phosphorylation (Oxphos) pathways to provide energy for various physiological activities, such as activation and proliferation[47, 48]. We found that the glycolysis of □*Mtb*-activated NKcells is significantly reduced, and this may explain the decline of IFN-□ production due to the growing consensus that glycolysis is critical for IFN- production by NK cells [49].

Immune cell metabolism plays a vital role in shaping immune responses to pathogen infection. Effector immune cells are believed to upregulate glycolysis to enable a quicker turnover of ATP, essentially switching to a state of aerobic glycolysis to meet the urgent demand for mounting response to pathogenic challenges in the form of increased proliferation, production of cytokines, and other cytotoxic capabilities[50]. This enhancement usually occurs through the upregulation of glycolytic enzymes and through the upregulation of surface nutrient transporters such as CD71, CD98, Glut1[51]. Several transcription factors are also involved in orchestrating the metabolic rewiring, NF-κ B (nuclear factor kappa-light-chain-enhancer of activated B cells), HIF-1α), (hypoxia-inducible factor-1α),c-Myc, Akt and mTOR (mechanistic target of rapamycin) are all known to differentially regulate the glycolytic gene expression landscape upon stimulation [52]. Apart from being a quicker source of energy, glycolysis also fuels the Pentose phosphate pathway, which increases the availability of PPP intermediates (ribose-5-phosphate and NADPH) essential for proliferation and effector functions [53]. Conversely, glycolytic end products can also be shunted into the TCA cycle as acetyl-CoA, NADH and FADH2 to further support OXPHOS, essentially supporting an energetic phenotype[54]. Recent studies have shown that NK cell lacking in Lactate dehydrogenase A loses their tumorigenicity and anti-viral function, suggesting an indispensable role of glycolysis[55–57].

Collectively, we conclude the mechanism that ALA is involved in as follows: ALA can function as a downregulator of NK cell glycolysis. The decreased glycolysis will then result in NK cell autophagy and the reduction of IFN- □production. We hypothesize that HIV can upregulate ALA and thus stimulate *Mtb* growth during HIV-*Mtb* co-infection. Our study offers a new understanding of the HIV-*Mtb* interaction and provides the implication of nutrition intervention and therapy for HIV-*Mtb* co-infected patients. This will be further investigated in our future work.

## Acknowledgments

We thank Dr. Raju S. R. Adduri for the metabolic analysis and for the editing of the manuscript.

## Author contributions

G.Y.: Conceived and designed research, analyzed data, performed experiments, drafted manuscript, edited, and revised manuscript.

B.Y.: Performed experiments and analyzed data.

T.M.: Performed experiments, analyzed data, and drafted manuscript.

R.R.: Performed experiments, analyzed data, and edited manuscript.

P.P.: Performed experiments and analyzed data.

D.A.: Performed experiments.

R.V.: Conceived, designed research, and edited manuscript.

D.T.: Conceived and designed research, analyzed data, and edited manuscript.

