## Supplementary figures for "HIV-differentiated metabolite N-Acetyl-L-Alanine dysregulates human natural killer cell responses to *Mycobacterium tuberculosis* infection"

**Supplementary Figure 1:**

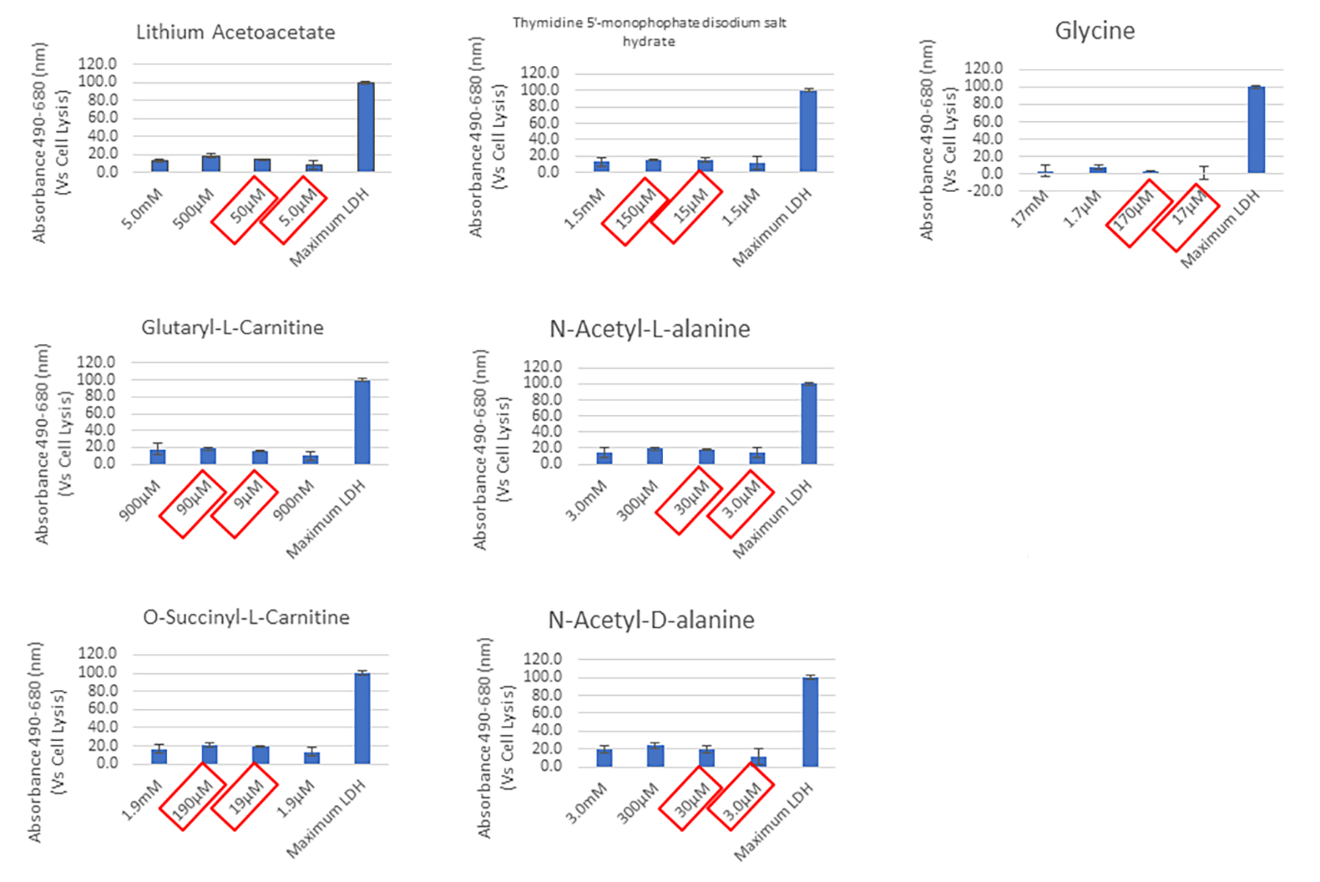

**Figure S1.** LDH assay was performed to determine the cytotoxicity of metabolic chemicals as described in the Methods section. The cytotoxicity was calculated as the percentage of positive control (the LDH assay kit provided) of the absorbance. For each chemical, two concentrations were chosen (shown in red boxes) for the following experiment shown in Figure 1.

**Supplementary Figure 2:**

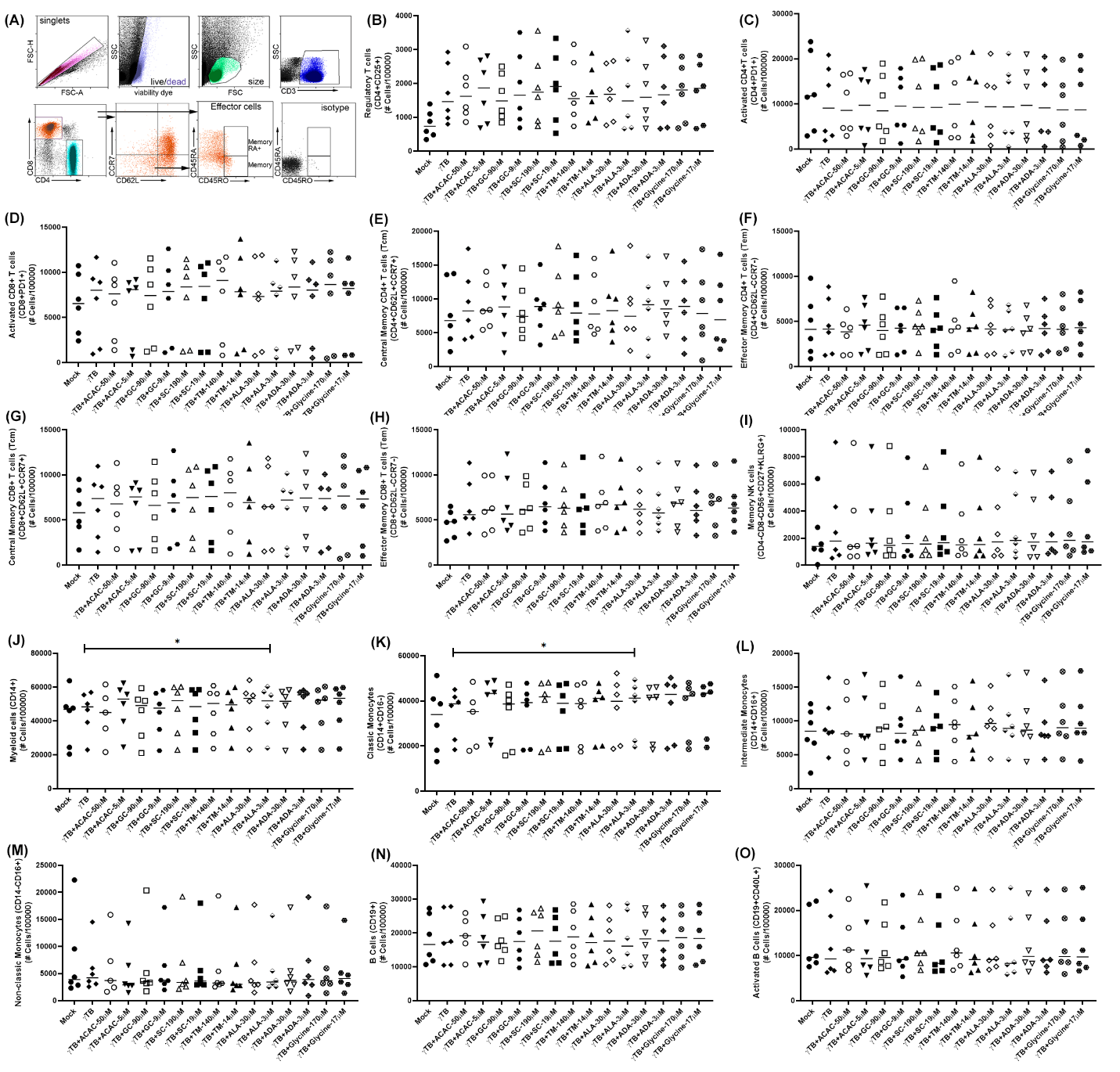

**Figure S2.** The effects of various metabolites on the expansion of various immune cells and their subpopulations. **(A)** Four panels were designed to stain the immune cells, and one typical panel is shown to describe the gating strategy of various phenotypes of T cells. **(B to O)** Statistical analyses of various immune cells and their subpopulations under different treatments. The paired T-tests were used to determine the differences between the treatment groups. The significant P values are shown (p<0.05).

**Supplementary Figure 3:**

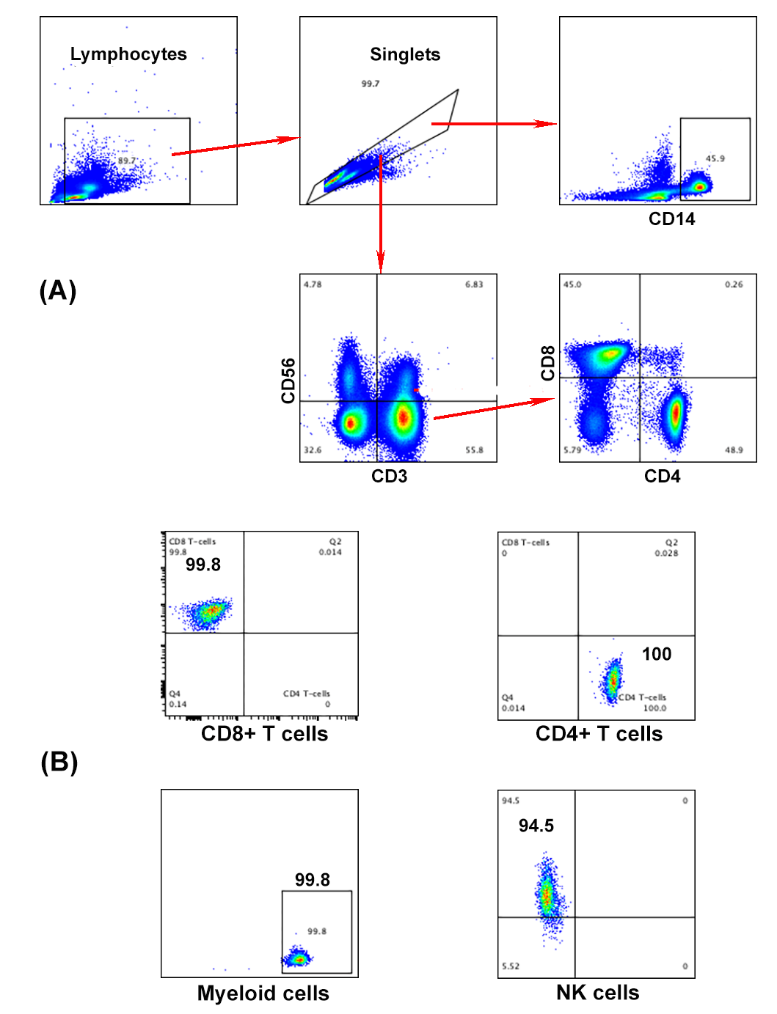

**Figure S3.** PBMCs from the LTBI+ donors were subjected to cell sorting to collect the CD4+ and CD8+ T cells, myeloid cells, and NK cells. **(A)** The gating strategy for cell sorting. **(B)** The purity of the collected CD4+ and CD8+ T cells, myeloid cells, and NK cells.

**Table S1:** Primer sequences used in Figure 5.

| **Name** | **Sequence** |
| --- | --- |
| IL21R-F | CTCGTCTGCTACACCGATTAC |
| IL21R-R | CTCGTCCTTCAGCTCTTCATAC |
| NKp46-F | TACTGCTCAAGGAGGGAAGA |
| NKp46-R | AGACCAGGCATGGTTGTTATAG |
| NKG2D-F | CAGCAAAGAGGACCAGGATTTA |
| NKG2D-R | GTTAGTAGGTTGGGTGAGAGAATG |
| KLRC1-F | GATTTACCATCAGCTCCAGAGAA |
| KLRC1-R | GTTACCACAGAGGCCATTAAGA |
| KLRG1-F | AGAGACTCACACCTCCTTGT |
| KLRG1-R | CCTCAGACCAATCCAGCAAA |
| KLRD1-F | GTACCGGTGCAACTGTTACT |
| KLRD1-R | CTGAAGCAGGCTGGATTTCT |
| DNAM1-F | AGGAGAAGGAGAGAGAGAAGAG |
| DNAM1-R | GATTGGTAGGTTGACTGGTAGAG |
| 2B4-F | CAGCATACAGACGAAGGTTGA |
| 2B4-R | GAGCCATTCTCCCACTTCAATA |
| CD27-F | TCAGTGTGATCCTTGCATACC |
| CD27-R | CGAACGAGAAGACCAGAGTTAC |
| CD16-1F | GACAATTCCACACAGTGGTTTC |
| CD16-1R | TTTGTCTGGCACCTGTACTC |
| Ki67-F | TAACACCATCAGCAGGGAAAG |
| Ki67-R | CTGCACTGGAGTTCCCATAAA |
| PRF1-F | CCTGTGAGGAGAAGAAGAAGAAG |
| PRF1-R | TCGTTAATGGAGGTGTGATGG |
| GZMA-F | GAGACTCGTGCAATGGAGATT |
| GZMA-R | CGAGGGTCTCCGCATTTATT |
| GZMB-F | ACACTCACACACACTACAAGAG |
| GZMB-R | ACGCACAACTCAATGGTACT |
| Tbet-F | AAGGATTCCGGGAGAACTTTG |
| Tbet-R | GTTGGGTAGGAGAGGAGAGTAG |
| Eomes-1F | GTGGCAAAGCCGACAATAAC |
| Eomes-1R | CCGAATGAAATCTCCTGTCTCA |
| STAT4-F | GCTTGGGCATCCATCATTTG |
| STAT4-R | GTGGCAGGTGGAGGATTATT |
| NFKB-F | CTCCACAAGGCAGCAAATAGA |
| NFKB-R | ACTGGTCAGAGACTCGGTAAA |
| AP1-F | TTCTATGACGATGCCCTCAAC |
| AP1-R | TCAGGGTCATGCTCTGTTTC |
| NFAT2-F | AATTCTCTGGTGGTTGAGATCC |
| NFAT2-R | TACTGGCTTCGCTTTCTCTTC |
| NFAT1-F | AGTGGCAGAATCGTCTCTTTAC |
| NFAT1-R | CAGCTGTCTGTGTCTTGTCTT |
| NFAT5-1F | CGTGGCTCAGTGAAAGATAGAA |
| NFAT5-1R | AGTCGTTGCCCACAAACA |
| **ActB-F** | CACTCTTCCAGCCTTCCTTC |
| **ActB-R** | GTACAGGTCTTTGCGGATGT |

**Table S2:** Primer sequences used in Figure 6.

| CASP1-F | GCTGAGGTTGACATCACAGGCA |
| --- | --- |
| CASP1-R | TGCTGTCAGAGGTCTTGTGCTC |
| CASP3-F | GGAAGCGAATCAATGGACTCTGG |
| CASP3-R | GCATCGACATCTGTACCAGACC |
| CASP8-F | AGAAGAGGGTCATCCTGGGAGA |
| CASP8-R | TCAGGACTTCCTTCAAGGCTGC |
| Bcl2-F | ATCGCCCTGTGGATGACTGAGT |
| Bcl2-R | GCCAGGAGAAATCAAACAGAGGC |
| MLKL-F | TCACACTTGGCAAGCGCATGGT |
| MLKL-R | GTAGCCTTGAGTTACCAGGAAGT |
| RIPK1-F | TATCCCAGTGCCTGAGACCAAC |
| RIPK1-R | GTAGGCTCCAATCTGAATGCCAG |
| IL1B-F | CCACAGACCTTCCAGGAGAATG |
| IL1B-R | GTGCAGTTCAGTGATCGTACAGG |
| TNR1-F | CCGCTTCAGAAAACCTCAG |
| TNR1-R | ATGCCGGTACTGGTTCTTCCTG |
| Atg3-F | ACTGATGCTGGCGGTGAAGATG |
| Atg3-R | GTGCTCAACTGTTAAAGGCTGCC |
| LC3B-F | GAGAAGCAGCTTCCTGTTCTGG |
| LC3B-R | GTGTCCGTTCACCAACAGGAAG |
| GPX4-F | ACAAGAACGGCTGCGTGGTGAA |
| GPX4-R | GCCACACACTTGTGGAGCTAGA |
| **ActB-F** | CACTCTTCCAGCCTTCCTTC |
| **ActB-R** | GTACAGGTCTTTGCGGATGT |
